# The GspLM inner membrane complex from the bacterial type II secretion system is a dimer of dimers and interacts with the system ATPase with high affinity

**DOI:** 10.1101/2020.03.20.999888

**Authors:** Aleksandra Fulara, Ioanna Ramou, Savvas N. Savvides

## Abstract

The type II secretion system (T2SS) is a multiprotein machinery spanning the diderm of gram-negative bacteria. T2SS contributes to the virulence of numerous gram-negative pathogens, including the multidrug resistant species *Pseudomonas aeruginosa, Acinetobacter baumanii, Klebsiella pneumonia* and *Vibrio cholerae*. Even though the T2SS has been studied extensively over the past three decades, our understanding of the molecular basis of its biogenesis and of its overall structure still remains unclear. Here we show that the core component of the inner membrane platform, the GspLM membrane protein complex, can be isolated as a dimer of dimers. Importantly, the complex is able to bind the T2SS ATPase, GspE, with high affinity. Finally, we have developed single domain VHH camelid antibodies (nanobodies) against the GspLM complex and have identified a nanobody that effectively prevents the cytoplasmic domain of GspL, GspL^cyto^, from binding to GspE. Our findings suggest that the T2SS ATPase is permanently associated with the inner membrane platform and that the GspELM complex should be considered as a key subassembly for the biogenesis of the T2SS apparatus.

## INTRODUCTION

The ability of bacteria to secrete proteins is essential for their communication with their external environment and for maintaining physiological processes such as adhesion, adaptation, and survival. For this purpose, gram-negative bacteria have developed at least seven secretion systems [1–3] of which the type II secretion system (T2SS) is best adapted for secretion of multiple exoproteins [4–7]. The system plays a role in the virulence of numerous human pathogens such as *Pseudomonas aeruginosa, Vibrio cholerae, Klebsiella pneumonia*, enterohemorrahagic *Escherichia coli, Burkhodelia cepacia* or *Leginella pneumophia* [5,8–10]. Currently, most of these pathogens imprint a serious threat to human health worldwide due to their resistance to antibiotics [11,12]. Therefore, understanding of bacterial virulence-related processes such as protein secretion is crucial for the development of much-needed novel treatment options against bacterial infections. Among gram-negative pathogens causing nosocomial infections, *P. aeruginosa* is a bacterium notoriously known for its efficiency in developing resistance to antibiotics [13,14].

T2SS is a nanomachinery spanning the diderm of gram-negative bacteria that facilitates secretion of folded proteins through the outer membrane [3,8,15]. In the case of *P. aeruginosa*, the system consists of 12 proteins, historically termed with the prefix Xcp. Due to its well-established homology with other T2SSs and the fact that the *Pseudomonas* species are the only terminology outliers, here we use the Gsp nomenclature to denote the T2SS proteins. The nexus of the system, the inner membrane platform (IMP), is composed of four integral membrane proteins: GspL, GspM, GspC and GspF. The components of the IMP interact with all the other subassemblies of the system: cytoplasmic ATPase GspE, periplasmic pseudopilus consisting of GspG, GspH, GspI, GspJ and GspK and the outer membrane secretin GspD. In addition, functional pseudopilins depend on the presence of prepilin peptidase GspO. Altogether, the four subassemblies form a functional secretion machinery that depending on species secretes proteins of diverse functions and destinations [7].

The IMP components GspL and GspM are bitopic proteins that interact with each other *in vivo* [16–19], and with the ensuing GspLM complex GspL is protected from proteolytic degradation [17,18]. Both GspL and GspM contain a single transmembrane helix followed by a periplasmic domain, and the periplasmic domains adopt the same rare permutation of the ferredoxin fold [20–22]. This similarity has been used as the basis for their proposed common evolutionary origin [21]. In contrast to GspM that only exposes a ~30-residue-long N-terminus to the cytoplasm, GspL possesses an extensive cytoplasmic domain that binds the system ATPase GspE [9,23–25] and interacts with the polytopic component of the IMP, GspF [26,27]. In addition, GspL can be crosslinked to the main pseudopilin GspG *in vivo*, which has been used as an argument for its role in coupling ATP hydrolysis to pseudopilus assembly [28]. GspM on the other hand has been suggested to play a role in aligning the IMP with the secretin channel based on the formation of discrete foci by fluorescently labeled GspM and GspC in the presence of secretin [29]. Recently, GspM has been shown to participate in the recruitment of the major pseudopilin to the membrane [30], and periplasmic domains of GspL and GspM were suggested to play a role in cargo protein recruitment [31].

Even though the individual proteins and domains composing the GspLM complex have been characterized, the GspLM complex has been barely studied as a whole. It is believed to be a dimer mostly based on oligomerization tendencies of soluble domains of GspM and GspL [20,21,24,32]. As we have shown recently, the periplasmic domain of GspL, GspL^peri^, adopts two oligomeric forms: a dimer and a tetramer [22]. Although in the context of an isolated domain the dimer of GspL^peri^ is a more stable oligomeric form, the influence of GspM on oligomeric tendencies of the GspLM complex remains an open question. In addition, this study is also devoted to the interaction at the base of the T2SS, the process of anchoring the system ATPase GspE in the inner membrane by GspL. Determination of kinetic parameters and affinity of the GspEL^cyto^ complex formation enables us to revisit the T2SS biogenesis scenario, according to which the ATPase is recruited to the inner membrane platform.

## RESULTS

### Recombinant production and quality assessment of the GspLM complex

GspL and GspM form the GspLM complex at the bacterial inner membrane and the complex is essential for the assembly and functionality of the IMP of T2SS [15,16]. Consistent with this functional role, the corresponding genes are adjacent in the *gspEM* operon [33]. Thus, we cloned an uninterrupted stretch of DNA to generate an expression vector that would enable co-expression of the full-length proteins and purification of the complex via strep-tag fused to C-terminus of GspM (**Figure 1a** and **1b**). Indeed, following detergent solubilization of GspLM we were able to purify the complex by affinity chromatography on a Strep-tactin column. The GspLM complex eluted as a single, monodisperse peak in the subsequent size exclusion chromatography purification **(Fig. 1c)**, and yielded 5 mg of pure GspLM complex from 10 liters of bacterial culture suitable for biochemical and biophysical studies.

**Figure 1.**
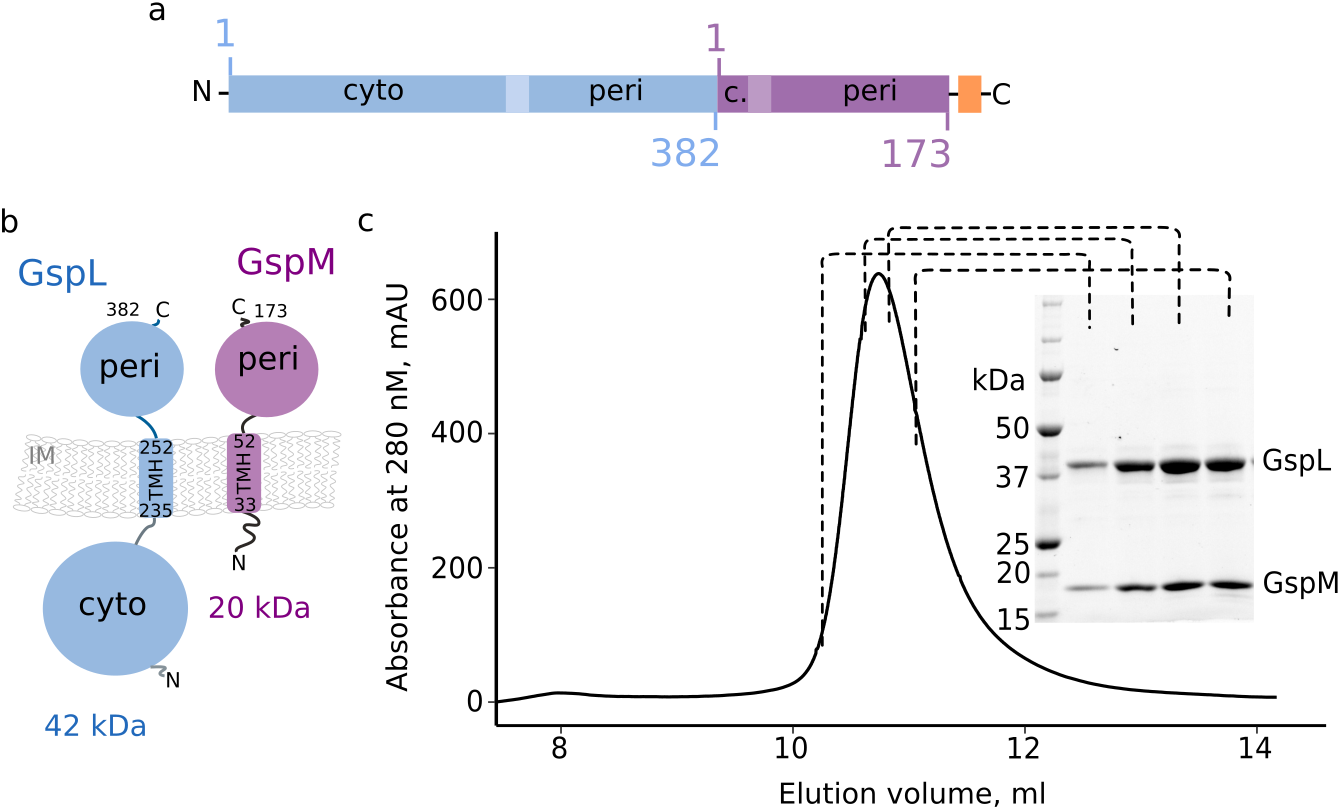
The GpLM complex: construct design, topology and elution profile. **(a)** The particular organization of the *gsp* operon enables the construction of an expression vector suitable for the co-expression of GspL and GspM. “Cyto” and “peri” refer to cytoplamic and periplamic domains, transmembrane helices (TMH) are marked with lighter colors and the strep-tag is marked in orange. The numbering applies to UniProt entries, indicated in Table 1. The domain and terminus annotation applies also to the topology scheme in **(b)**. IM: inner membrane, **(c)** Size-exclusion chromatogram of recombinant GspLM and the corresponding SDS-PAGE analysis. The GspLM complex elutes as a single peak and the molecular weights (MWs) of the complex components resolved on the gel are in good agreement with the theoretical MWs (indicated in panel b).

### GspLM is a dimer of dimers in solution

Understanding the instrinsic oligomeric states of protein subassemblies of the bacterial T2SS is necessary for deciphering the architectures and assembly principles of the functional T2SS apparatus. With the exception of the outer membrane secretin [34–36], experimental data on the oligomeric state of the membrane-embedded components of the T2SS is scarce. Therefore, we addressed the question of the oligomeric state of the purified GspLM complex via SEC-MALLS. This non-invasive biophysical method allows an accurate mass estimation of protein:detergent micelle conjugates.

**Figure 2a** shows that the GspLM complex elutes as a single peak corresponding to a protein mass of 133 kDa and a detergent micelle of 48 kDa. Since the theoretical molecular weight of the complex calculated from the sequence is 61.5 kDa, the experimental molecular weight can be readily explained by a dimer of dimers. In order to delineate the contributions of individual proteins to the GspLM dimerization interface, we subjected their soluble domains namely GspM^peri^, GspL^peri^, GspL^cyto^ to SECMALLS. Our analyses show that GspM^peri^ and GspL^cyto^ are monomers, whereas GspL^peri^ is a dimer in solution **(Fig. 2b-d)**. Based on this result, it is more likely that GspL^peri^ participates in the GspLM dimer interface rather than GspL^cyto^ or GspM^peri^. The intrinsic dimerization propensity of GspL^peri^ was expected, as its dimeric interface has recently been shown to feature extensive hydrophobic interactions [21,22], leading to obligate GspL^peri^ dimers in solution.

**Figure 2.**
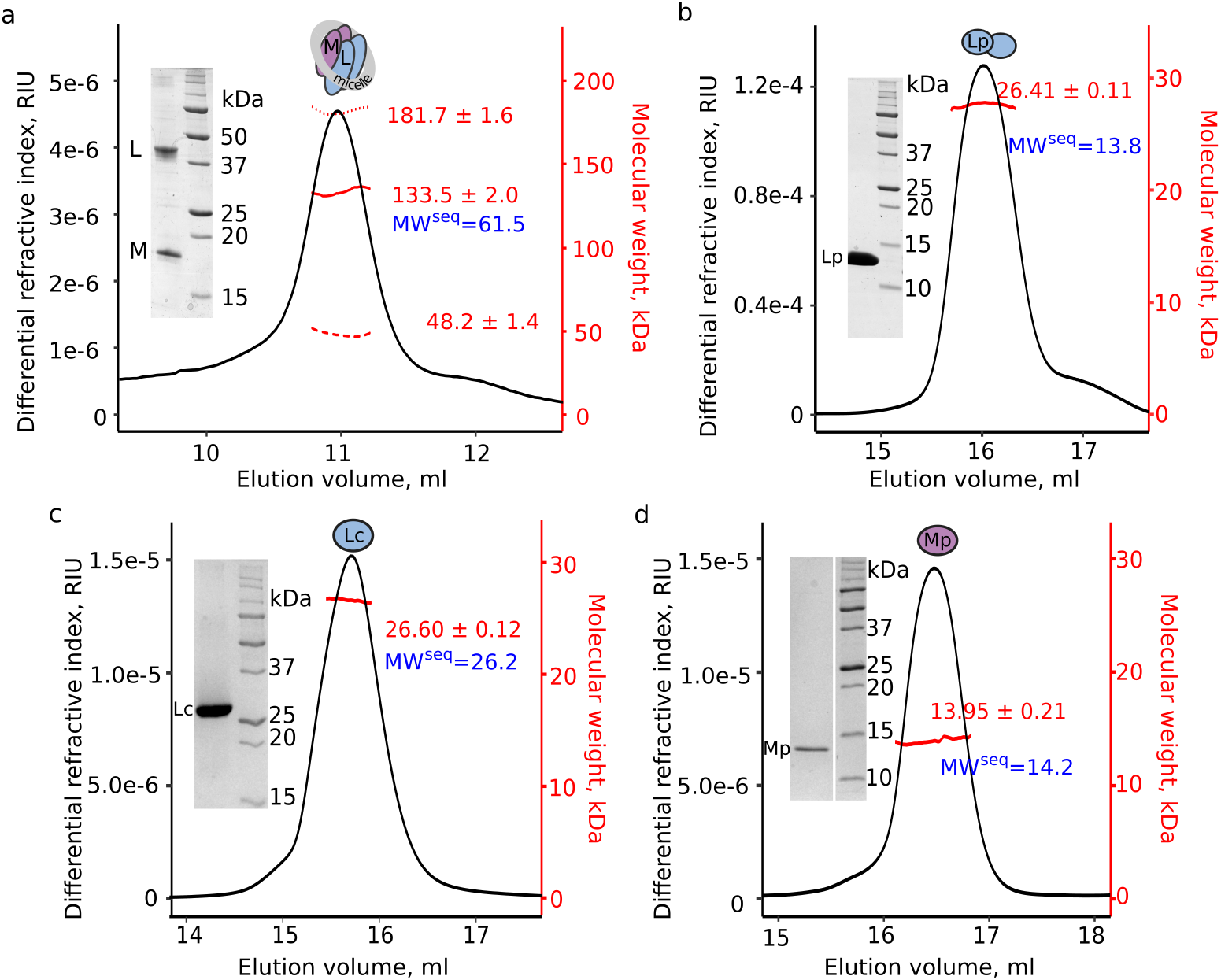
GspLM is a dimer of dimers. SEC-MALLS analysis of **(a)** the GspLM complex and its soluble domains: **(b)** GspL^peri^, **(c)** GspL^cyto^, and **(d)** GspM^peri^. The experimentally determined molecular weight is plotted across the chromatographic peak and is reported in kDa ± standard deviation in red, whereas the monomeric masses calculated from the sequences are in blue. In case of GspLM the detergent contribution is distinguished. Red solid line: protein mass, red dotted line: total mass, red dashed line: detergent mass. Insets contain SDS-PAGE analysis of samples subjected to SEC-MALLS. On the gels, the experimental samples (left lines) are accompanied with molecular weight marker (right lines, in kDa). The theoretical MWs of monomers are in a good aggreement with masses derived from SDS-PAGE analysis..

However, we were surprised with the finding that neither GspL^cyto^ nor GspM^peri^, which were previously shown to possess dimerization tendencies [19], do not retain this oligomeric state in their isolated forms. Thus, conclusions regarding the physiological context of such oligomeric forms can only be relevant in light of full-length complexes. On the other hand, relevance of the oligomeric state of the complex has to be always cross-validated while working with *in vitro* preparations. In this regard, in the following experiments we employed the cytoplasmic interaction partner of GspL, the system ATPase GspE.

### Recombinant GspLM interacts with the system ATPase GspE with high affinity

GspL and GspM have no enzymatic activity and therefore their ability to interact with each other and with other Gsp proteins might serve as a way to evaluate their functional competence. The fact that GspL and GspM copurify to form a stable complex with 2:2 stoichiometry **(Fig. 1c and 2a)** represents a first argument for the relevance of the complex. In the sequence of protein-protein interactions leading to the secretion of a cargo protein, GspL is thought to anchor the system ATPase GspE and to transduce the energy from ATP hydrolysis to the periplasm [9,24,25,37]. Therefore, the possibility to examine the GspL – GspE interaction by using GspL embedded in the GspLM complex appears as a venue to test the functionality of the recombinant GspLM complex.

We probed the GspE – GspLM via BLI using the full-length GspE as a ligand and detergent-solubilized full-length GspLM complex as an analyte over a concentration range of 0.2-300 nM **(Figure 3a)**. The obtained binding curves illustrate that the GspLM complex interacts specifically with GspE to form a ternary GspELM complex with a binding affinity in the nanomolar range. On the other hand, the curves could not be be credibly fitted with any available kinetic model, which prevented us from obtaining a quantitative characteristics of the ternary complex formation. We assume that the detected GspL-GspE interaction, that is mostly driven by true chemical affinity, might be contaminated by a low degree of aspecific interactions. We are aware that a common way to prevent such scenario in BLI experiments is to supplement the reaction buffer with a nonionic detergent, Tween-20. Unfortunately, this detergent is detrimental to the stability of the GspLM complex. Alternative supplementation of the BLI reaction buffer with nonionic detergents that do not compromise integrity of GspLM did not improve the shape of the curves. In order to bypass the difficulty of working with the membrane protein complex and given the fact that the cytoplasmic domain of GspL, GspL^cyto^, interacts with GspE [24,37], GspL^cyto^ was used as an analyte in the following experiments.

**Figure 3.**
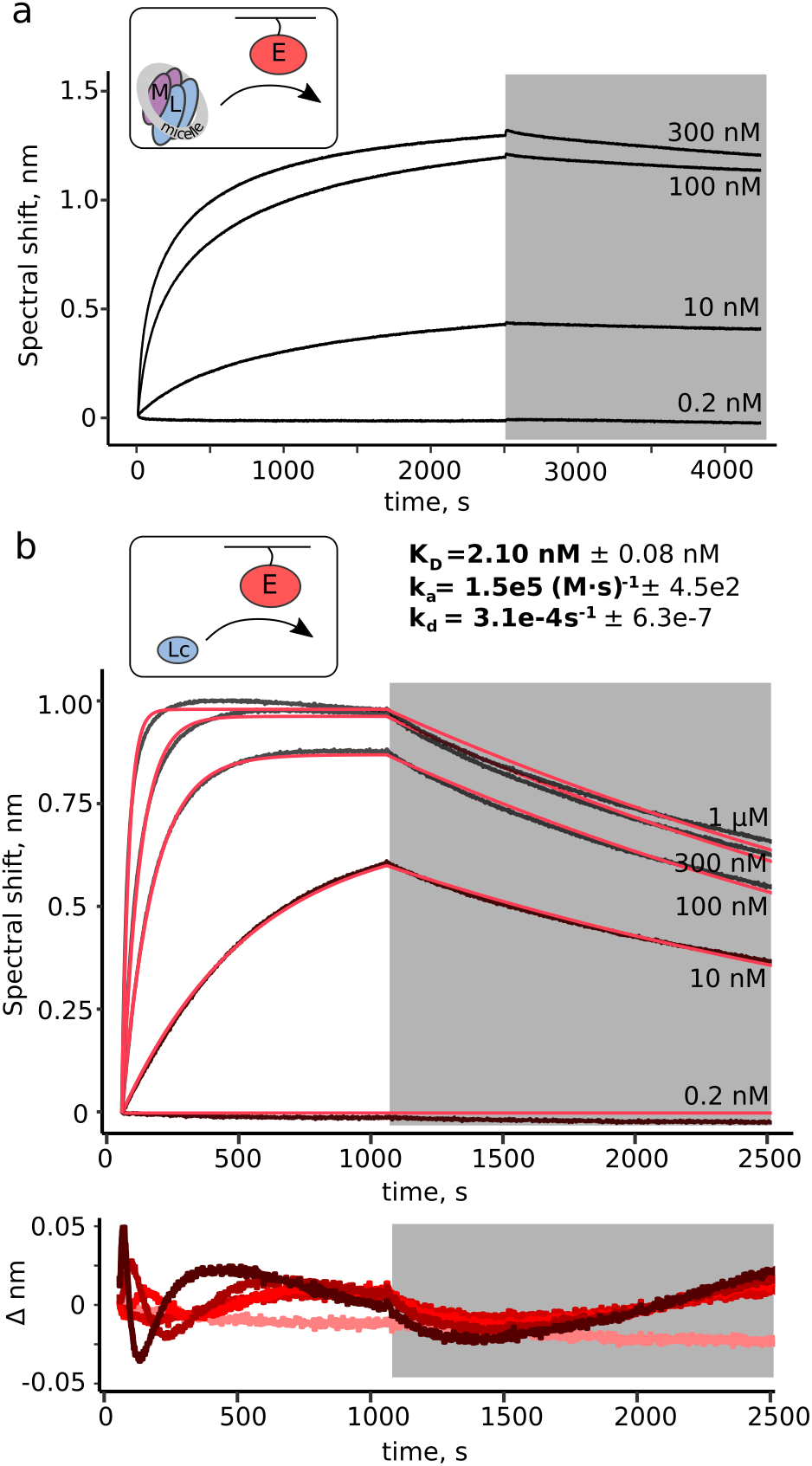
GspL and GspE interact with high affinity. **(a)** The GspELM and **(b)** the GspEL^cyto^ complex formation followed with BLI. In each case the biotinylated GspE was used as ligand. Analyte concentrations are indicated next to each trace. Insets contain schematic representation of each experiment. The grey ring arround GspLM represents detergent micelle. E: GspE, LM: GspLM, Lc: GspL^cyto^. In experiment (b) the data were fitted with a 1:1 binding model (in red). Residual trace of the fit is presented in the bottom panel. Color of the trace darkens with the increase of concentration. The reported K_D_, k_a_ and k_d_ values represent average values ± standard deviation.

### GspL^cyto^ and GspE interact with high affinity

The kinetic analysis of the GspE-GspL^cyto^ interaction is presented in **Figure 3b**. The binding event is characterized by the kinetic constants *k_a_* and *k_d_*, which lead to a dissociation constant *K_D_* of 2.1 nM. The affinity of the GspE-GspL^cyto^ interaction is reported here for the first time and the nanomolar range enables to classify it as a high affinity interaction. In this regard, the GspE-GspL^cyto^ interaction appears to be rather permanent, in comparison to the transient associations of GspC-GspD and the interactions between a cargo protein LasB and the multiple periplasmic domains of Gsp proteins [38–40]. Thus, the herein determined nanomolar affinity of the GspE-GspL^cyto^ interaction provides an argument to discuss biogenesis of the T2SS.

### Development and characterization of VHH single domain antibodies against GspLM

In order to facilitate further characterization of the GspLM complex, a library of VHH single domain camelid antibodies, known as nanobodies (nb), was generated. A nanobody consists of a single monomeric variable domain of a heavy-chain only antibody, of 12-15 kDa. Their high antigen specificity and low molecular weight together with ease of expression and purification make such antibody fragments very well suited for the structural characterization of membrane proteins and protein complexes [35,41–43]. Following immunizations with the GspLM complex we obtained eight nanobody classes and opted to determine the binding epitopes of representatives from each class. Furthermore, we were primarily interested in identifying nanobodies that would target the interface of GspL and GspM in the GspLM complex.

The binding capacity of the selected nanobodies has been tested with BLI, as presented in **Figure 4**. All nanobodies except for nb8 bind the GspLM complex with nanomolar affinity **(Fig. 4a)**. The tested nb8 was the only member of class 8, and therefore it could not be replaced **(Supporting Figure S1)**. Further on, we sought to determine binding epitopes of the seven remaining nanobodies. For this reason the BLI sensors were loaded with nanobodies and subsequently immersed in concentrated solutions of GspL^peri^, GspM^peri^ and GspL^cyto^ (see *Methods* for details). Four nanobodies out of seven, namely nb2, nb4, nb5 and nb6, bind the cytoplasmic domain of GspL **(Fig. 4b)**. Interestingly, the three remaining nanobodies, representatives of classes 1, 3 and 7, do not bind either GspL^peri^, GspL^cyto^ or GspM^peri^ **(Fig. 4b)**. Nevertheless, they all interact with the GspLM complex **(Fig. 4a)**. In this regard it is of interest to assign the binding epitope of nanobodies 1, 3, and 7.

**Figure 4.**
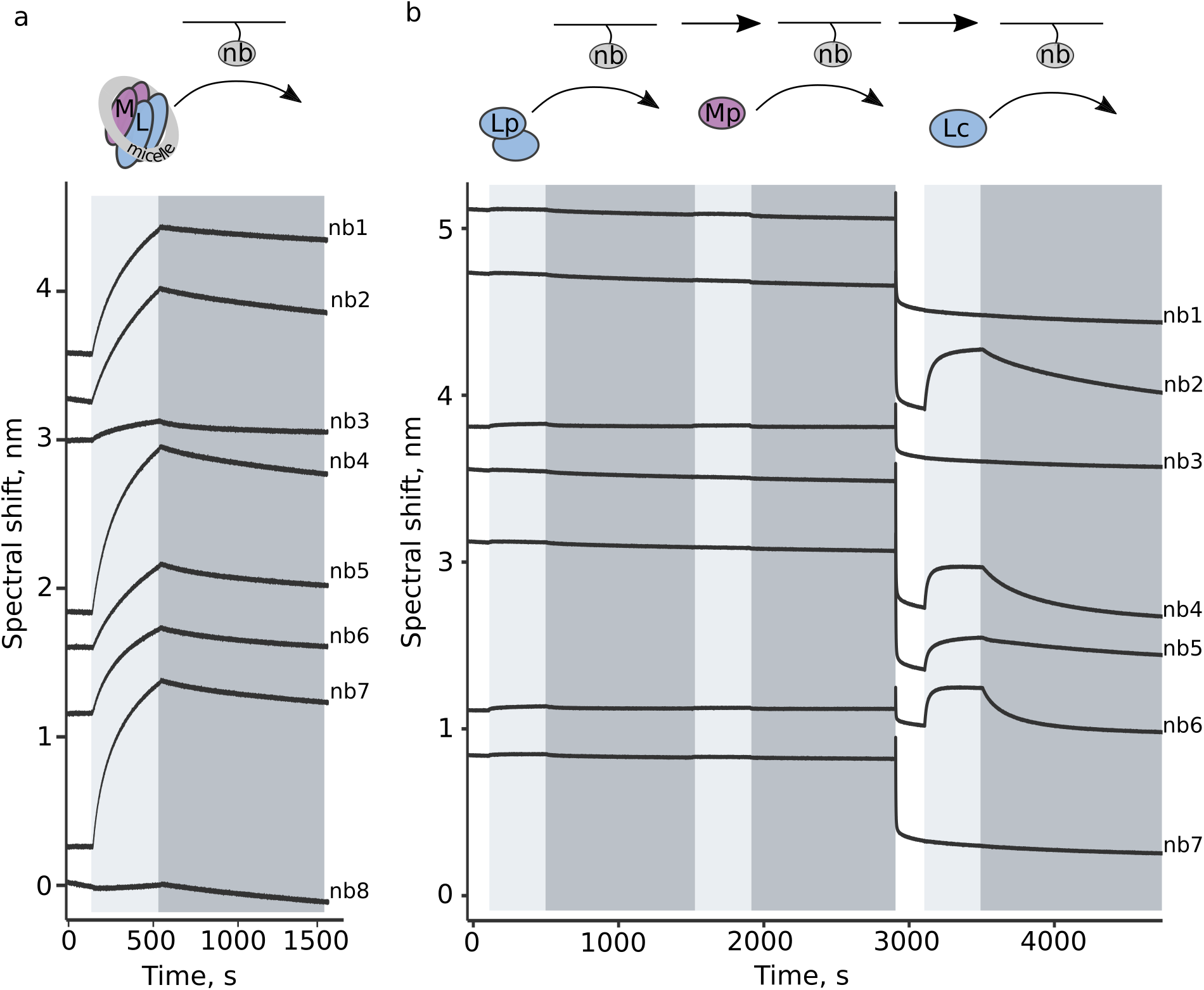
Delineation of the binding epitopes of the anti-GspLM nanobodies. Upper panels contain schematic representations of each experiment and bottom panels contain BLI traces of formation of complexes. **(a)** Nanobody (nb) representatives of classes 1-7 bind the GspLM complex, **(b)** Nanobodies 2, 4, 5 and 6 bind the cytoplasmic domain of GspL. Association and dissociation phases are highlighted in light and dark grey, respectively. LM: GspLM, Lp: GspL^peri^ Mp: GspM^peri^, Lc: GspL^cyto^ The grey ring arround GspLM represents detergent micelle.

GspLM consists of the three tested soluble domains, in addition to the transmembrane helices (TMHs) and the ~30 residue long N-terminus of GspM, which is exposed to the cytoplasm **(Fig. 1b)**. The TMHs, which were either surrounded by the amphipol belt (as for the immunizations for nanobody generation, see *Methods* for details) or embedded in a detergent micelle (as in the experiment presented in **Fig. 4a**), were sterically unavailable as nanobody binding epitopes. The N-terminus of GspM is more likely to be immunogenic than the TMHs, although its close proximity to the membrane might result in limited exposure. On the other hand, formation of the GspLM complex results in a novel structural assembly, which as such could trigger the generation of antibodies specific to the complex rather than the individual protein components. It is therefore probable that nanobodies 1, 3 and 7 might engage a binding epitope at the GspLM interaction interface.

### nb2 is a GspL^cyto^ antagonist for its interaction with the ATPase GspE

The availability of four GspL^cyto^ binders **(Fig. 4b)** prompted us to examine their ability to interfere with the high affinity interaction between GspL^cyto^ and the ATPase **(Fig. 3)**. To this end, GspL^cyto^ was first saturated with ~20-fold excess of nanobodies, as compared to respective affinities **(Table S2)**. In a second step, the interacting proteins were crosslinked, to prevent dissociation. In the last step, the GspL^cyto^-nb complexes were tested for binding against recombinant GspE (**Fig. 5a**, see *Methods* for details). We reasoned that if a nanobody competes with GspE for the same epitope, the BLI response elicited upon GspL^cyto^-GspE binding will be lower in the presence than in the absence of the nanobody.

The experiment revealed that GspL^cyto^ binds GspE to a much lesser extent when nb2 is present **(Fig. 5a)**. Importantly, the effect is specific, because GspE mixed with a non-GspLM specific nanobody binds GspL^cyto^ to the same extent as GspE alone (**Fig. 5a**, C_E vs. C_nb traces). In order to confirm the antagonistic effect of nb2 on the GspE-GspL^cyto^ interaction another competition binding experiment was performed, in which the impact of nb2 on the affinity of the GspE-GspL^cyto^ complex formation was evaluated **(Fig. 5b)**. Whereas pure GspL^cyto^ binds GspE with a K_D_ of 2 nM, the interaction is completely abolished in the presence of nb2 **(Fig. 5b)**. The measured affinity is consistent with the value we reported in **Figure 3b**. In light of the above, nb2 appears to serve as an antagonist for GspL^cyto^ by effectively preventing its interaction with the ATPase GspE.

## DISCUSSION

The bacterial type II secretion system is a membrane embedded nanomachinery, important for the virulence of numerous gram-negative pathogens. The GspLM complex constitutes the core of the inner membrane platform of the T2SS and provides a physical link between the cytoplasmic energy source and the periplasmic stages of the secretion process. Even though it has been researched for three decades, many aspects of the functionality of the complex have remained enigmatic. Our studies shed light onto the oligomeric organization of the GspLM complex and complement the knowledge on its interaction with the system ATPase GspE.

Based on the survey of scientific literature, the oligomeric state of the GspLM complex has never been addressed directly. Rather, it has been discussed in light of the oligomeric state of the individual proteins. The full-length GspL and GspM proteins of *V. cholerae* and *D. dadantii* were shown to homodimerize [17,19], and the interfaces were proposed based on structural models of their soluble domains [9,20,21,32]. Furthermore, the periplasmic domain of GspL forms not only dimers, but also tetramers in solution [22]. In contrast, the homologous PilNO complex from the T4PS of *P. aeruginosa* has been characterized recently, although in the absence of transmembrane segments of the proteins. The truncated PilNO complex [44] eluted from a size exclusion column at the size consistent with the 2:2 complex, containing two molecules of PilN and two molecules of PilO. Interestingly, the tendency of T2SS protein components to form dimers as a minimal oligomeric assembly has also been illustrated in the case of *P. aeruginosa* GspD and GspF [45,46].

The availability of pure and stable full-length GspLM complex allowed us to address its oligomeric state directly via SEC-MALLS. In addition, examination of oligomeric states of the soluble domains with the same technique enabled us to assess the possible contributions of the individual protein domains to the GspLM oligomeric interface.

Our systematic SEC-MALLS study has revealed that the GspLM complex is a dimer of dimers **(Fig. 2a)** and that GspL^peri^ is its only purified soluble component that dimerizes on its own **(Fig. 2b-d)**. Based on the data presented in **Figure 2** and the hydrophobic nature of the GspL^peri^ dimer [22] one would expect that the oligomerization of GspLM significantly relies on the periplasmic domain of GspL. Importantly, the biochemically isolated complex retains its ability to interact with the secretion ATPase GspE with high affinity **(Figure 3a)**, which illustrates its functional relevance.

Demonstration of the GspE-GspL interaction **(Figure 3a)** not only proved the relevance of the isolated GspLM complex, but also motivated us to study the ATPase – GspL interaction in more detail, namely to determine the affinity of complex formation. In recent years, a hypothesis has been formulated that the assembly of a functional T2SS apparatus might be transient, relying on a set of low affinity protein-protein interactions [3,30,47,48]. Low affinity interactions have been shown to drive periplasmic stages of the secretion process, namely exoprotein recruitment by GspC and GspC-GspD association [38–40]. Nevertheless, previous studies suggested that the GspL-GspE interaction might be of high affinity [9,24,37]. With our BLI-based experimental setup we wanted to test how the stage of cytoplasmic recruitment of the ATPase fits the narrative of the T2SS assemblage.

The experimentally determined high affinity of the GspEL^cyto^ complex formation as reflected by a K_D_ equal to 2 nM **(Figure 3b)** strongly suggests the permanent character of the interaction. It is tempting to propose that high affinity interactions between T2SS protein components might drive association of the subassemblies, whereas low affinity interactions might guide the secretion process, which requires the combination of the subassemblies. In models describing the T2SS mode of action, the ATPase is handled either altogether with the IMP or separately, as a subassembly in the cytoplasm. In the latter case a recruitment stage is distinguished, whereby GspL engages in the interaction with GspE and anchors it in the inner membrane [15,48]. Our findings support the notion that the ATPase is permanently associated with the IMP, rather than recruited to it. In this regard the IMP-ATPase complex should be considered as a single subassembly.

Identification of a nanobody that acts as a GspL^cyto^ antagonist in its interaction with ATPase GspE provides a key new tool for facilitating further characterization of the structure and mechanism of bacterial T2SS. Although the medical potential of nanobodies is well recognized [49,50], this particular antagonist has limited chances to be followed as a potential therapeutic, due to the cytoplasmic residency of its target.

With this work we deliver hitherto missing biophysical and biochemical details on the GspLM complex and its association with the T2SS ATPase GspE. We show that the GspLM complex can be isolated as a stable dimer of dimers and that the interaction between the inner membrane platform and the ATPase is of high affinity. These findings are of particular value and timely, at the dawn of holistic characterization of the type II secretion system (bioRxiv deposits: [51,52]). The limited resolution of the megadalton complexes require restraints to build models that are biologically sound and our work provides them in a form of an oligomeric state restrains. Additionally, determination of high affinity interaction at the base of the T2SS provides new arguments for the discussion on identities of the subassemblies and the system biogenesis. Altogether, we shed light onto the architecture and biogenesis of the T2SS, the long studied but still enigmatic aspects of this secretion machinery.

## METHODS

### Construct design

Constructs used in the study are listed in **Supporting Table 1**.

**Table 1.**
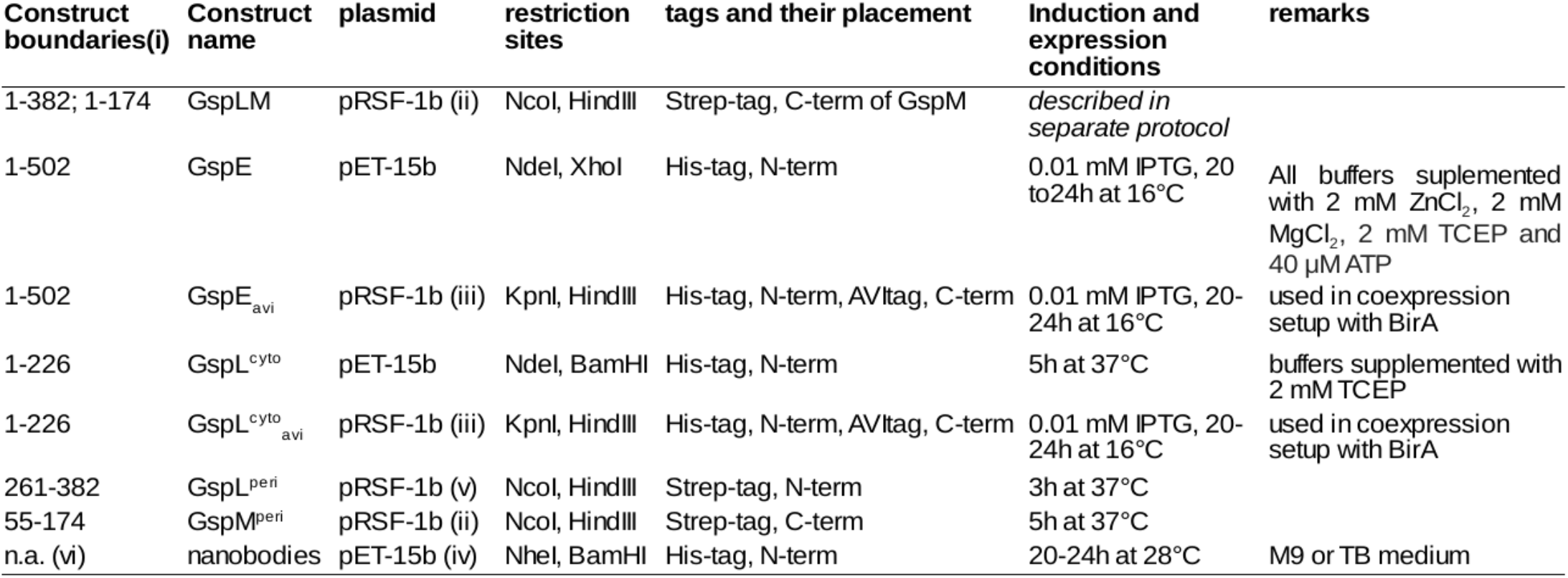
Constructs used in the study. The Table lists the differences between standard expression and purification protocols reported in Experimental procedures section. **(i)** applies to proteins with Uniprot codes: GspL: P25060, GspM: P25061, GspE: Q0512. **(ii)** StrepTAG introduced on a primer 5’-CCTAGAAGCTTTCATTATTTTTCGAACTGCGGGTGGCTCCAGCCGC TGCCGCCCTCGACCCGCAGGCTCAGG-3’ **(iii)** AVItag introduced on an oligo compatiibile with Hindill/Xhol restriction sites: 5’-AAGCTTGGAGGAGGATTGA ACGATATCTTCGAGGCACAAAAGATCGAGTGGCATGAGtaaCTCGAG-3’ **(iv)** pelB signal sequence followed by six histidine HisTAG was introduced on an oligo compatibile with Ncol and BamHI restriction sites: 5’-CCATGGGCAAATACCTATTGCCTACGGCAGCCGCTGGATTGTTATTACTCGCGGCCCAGCCGGCGATGGCCGCCCATCATCATCATCAT CATGATGAAGTTGÄTGGAGCTAGCGGAGAATCCGGATCC-3’ **(v)** StrepTAG introduced on a primer. 5’-AGCATCCATGGAATGGAGCCACCCGCAGTTCGAAAAACATATGA TGACAGCCCCATTACC-3’ **(vi)** n.a. not applicable.

### Expression of recombinant GspLM

*E. coli C43(DE3)* electrocompetent cells were transformed with DNA of GspLM in a pRSF1b expression vector backbone (**Table S1**) inoculated for 1h at 37°C and plated on an agar plate supplemented with kanamycin (Kan), 25 μg/ml. The next day a colony was picked up for inoculation in LB+Kan (lysogeny broth) and the culture was grown overnight. Afterwards the culture was used to inoculate an expression culture, in a 1:50 v/v ratio, in autoinduction (AI) medium. The cultures were grown for 3-4h at 37°C and afterwards the cells were transferred to 28°C for 20-24h. The AI medium, due to presence of phosphate buffer, was supplemented with higher amount of Kan (100 μg/ml).

### Membrane preparation and purification of recombinant GspLM

The frozen cell pellet was thawed and resuspended in a lysis buffer, 15ml per pellet from 1l culture (50 mM TRIS, pH=8.0 at 4°C, 300 mM NaCl, 20 mM βMe, HEWL: 1mg/g pellet) supplemented with Complete (Roche) EDTA-free protease inhibitor cocktail. The cells were lysed with pre-cooled Emulsiflex c3 homogenizer (Avestin), by two passages at 1.000 bar. For the purpose of membrane isolation, the homogenized mixture was centrifuged for 30 min at 15.000 g followed by 1h centrifugation at 100.000g. The pelleted membranes were homogenized in resuspension buffer, 1:1 v/v (50 mM TRIS, pH=8.0 at 4°C, 300 mM NaCl, 20 mM βMe) and frozen in 1 ml aliquots for long term storage.

Prior to purification, GspLM containing membranes were thawed, supplemented with dodecyl maltoside or decyl maltoside (DDM or DM) to a final detergent concentration of 3% and solubilized for 30 min with aid of a magnetic stirrer. The solubilization was followed by 30 min centrifugation at 100.000g. Up till this point all steps were performed at 4°C, whereas the following steps were done at room temperature. The supernatant was decanted and loaded on Strep-tactin Superflow matrix (IBA Lifesciences) equilibrated with a wash buffer (WB; 0.02% w/v DDM or 0.2% w/v DM, 50 mM TRIS, pH=8.0, 150 mM NaCl, 2 mM TCEP). The matrix-bound protein was washed with 15 column volumes (CV) of WB and eluted with 2xCV of WB containing 2,5 μM desthiobiotin (IBA Lifesciences). The eluate was concentrated and subjected to size exclusion chromatography on Superdex 200 increase semipreparative column (10/300, bead volume: 24 ml; GE Healthcare), also equilibrated with WB. The complex eluted in a single peak followed by a shoulder being an excess of GspM. Based on SDS-PAGE analysis, only the fractions containing the stoichiometric GspLM complex were pooled and concentrated for further experiments.

For the purpose of nanobody generation GspLM was transferred to amphipols, a surfactant that enables handling of the membrane protein in a detergent-free aqueous solution, by binding to the hydrophobic transmembrane region [53]. The size exclusion purified GspLM was concentrated and mixed with 10:1 molar excess of Amphipol A8-35 (Anatrace), for 30 min at 4°C. Afterwards, a 20:1 w/w excess of biobeads (Biorad), in relation to the amount of detergent, was added and the sample was incubated overnight at a magnetic stirrer at 4°C. The biobeads were removed by centrifugation at 20.000g for 30 min. The quality of the amphipol associated complex was controlled by performing an additional size exclusion step on a Superdex 200 increase column (10/300, bead volume: 24 ml; GE Healthcare).

### Standard protocol for expression and purification of soluble constructs

Numerous constructs were used in the studies, most often as purified proteins. The employed expression and purification protocols have multiple steps in common and such a standard protocol is reported here. If expression and/or purification of a particular construct deviates from this standard protocol, all the differences are listed in **Table S1**.

If not stated otherwise, a protein was expressed in the BL21(DE3) *E.coli* strain from an IPTG inducible plasmid using LB as a culture medium. The expression cultures were grown in shaker flasks at 37°C till OD=0.6 followed by induction with 1 mM IPTG. Expression timing and temperature conditions are listed in **Table S1**. The cells were harvested by 10 min centrifugation at 8.000g. At this point, the cell paste was stored at −80°C. When needed, the cell pellets were thawed and lysed by sonication in lysis buffer (50 mM TRIS, pH=8.0 at 4°C, 300 mM NaCl, HEWL: 1mg/g pellet) supplemented with Complete (Roche) EDTA-free protease inhibitor cocktail. Upon 30 min centrifugation at 75.000 g, the soluble fraction was filtered with a syringe cap filter (0.22 μm) and loaded on a proper matrix: either Ni-NTA Agarose (Qiagen) or Strep-tactin Superflow (IBA Lifesciences), depending on the purification tag exploited in the affinity step (indicated in **Table S1**). For nickel affinity chromatography, the filtered supernatant was loaded on the matrix equilibrated with Ni-NTA buffer (50 mM TRIS, pH=7.4, 500 mM NaCl, 25 mM imidazole). Afterwards the protein was washed with 10 CV of 50 mM imidazole Ni-NTA buffer and eluted with 2 CV of 300 mM Ni-NTA buffer, most of the time in batch mode. For Strep-tactin affinity chromatography the filtered supernatant containing the protein of interest was loaded on the matrix equilibrated with Strep-tactin buffer (50 mM TRIS, pH=7.4, 500 mM NaCl). The protein was washed with 10-15 CV of the Strep-tactin buffer and eluted with 2-3 CV of the Strep-tactin buffer supplemented with 2.5 μm desthiobiotin, most of the time in batch mode. As the final polishing step size exclusion chromatography was performed, employing Superdex 75 or Superdex 200 (depending on protein size). The standard SEC buffer was 50 mM TRIS, pH=8.0, 100 mM NaCl.

### SEC-MALLS

SEC-MALLS experiments were carried out on a Wyatt Technology system, consisting of a HPLC unit with a Superdex 200 increase column (10/300, GE Healthcare), UV detector (Shimadzu), a three-angle static light scattering detector (DAWN Theros) and a refractometer (Optilab T-rEX). The measurements were performed on purified protein samples at a concentration of 0.5-1.5 mg/ml, in buffers identical to the ones used in the purification step. Protein and DM dn/dc values of 0.185 ml/g and 0.147 ml/g respectively, were used for molecular weight calculation. Data analysis was performed with Astra V software, using the conjugate analysis module in the case of the membrane protein complex.

### Production of in vivo biotinylated proteins for BLI studies

GspE and GspL^cyto^ were cloned in the modified pRSF1b vector containing a C-terminal AVI-tag **(Table S1)**. Afterwards they were coexpressed with *E. coli* biotin ligase (BirA) on pET15b under conditions indicated in **Table S1**. Prior to BLI measurement a cell pellet aliquot originating from 10ml of expression culture was thawed, resuspended in 1 ml of kinetic buffer (50 mM TRIS, pH=8.0, 150 mM NaCl, 0.01% bovine serum albumine (BSA), 0.02% Tween-20), and lysed by sonication. Upon centrifugation the supernatant was diluted 5-20x in kinetic buffer and used for functionalization of streptavidin-coated tips. The proteins were further used in BLI experiments.

### BLI

The experiments were performed on an Octet RED96 instrument (ForteBio), operating at 25°C with employment of either streptavidin-(SA) or Ni-NTA-coated biosensors, which is indicated in the legends of the relevant figures. The composition of kinetic buffers is 50 mM TRIS pH=8.0, 150 mM NaCl, 0.01% bovine serum albumine (BSA), 0.05% Tween-20 for SA-based experiments and 50 mM HEPES pH = 7.4, 150 mM NaCl, 0.01% BSA, 0.05% Tween-20 for Ni-NTA-based experiments, with the exception of experiments whereby GspLM was an analyte. The aforementioned experiments were performed in 0.02% w/v DDM, 50 mM TRIS, pH=8.0, 150 mM NaCl, 2 mM TCEP. In all experiments the association step was preceded by reaching baseline signal in kinetic buffer. All experiments were performed at least in triplicates and the reported parameters are average values from these measurements.

The streptavidin-coated tips were functionalized with biotinylated GspE (**Figs. 3, 5b**) or GspL^cyto^ **(Fig. 5a)**, whereas Ni-NTA-coated tips were functionalized with N-terminally His-tagged nanobodies **(Fig. 4)**. The loading of His-tagged ligands was followed by a covalent linkage step being a 2 min dip in 10 mM 1-ethyl-3-(3-dimethylaminopropyl)carbodiimide hydrochloride (EDC), 20 mM N-hydroxysulfosuccinimide (Sulfo-NHS) solution and a quenching step being a 2 min dip in 1 M TRIS, pH=8.0. The covalent linkage/quenching tandem was also performed in the experiment presented in **Figure 5a**, after capturing the GspL^cyto^/nanobody interaction and before exposure to GspE.

**Figure 5.**
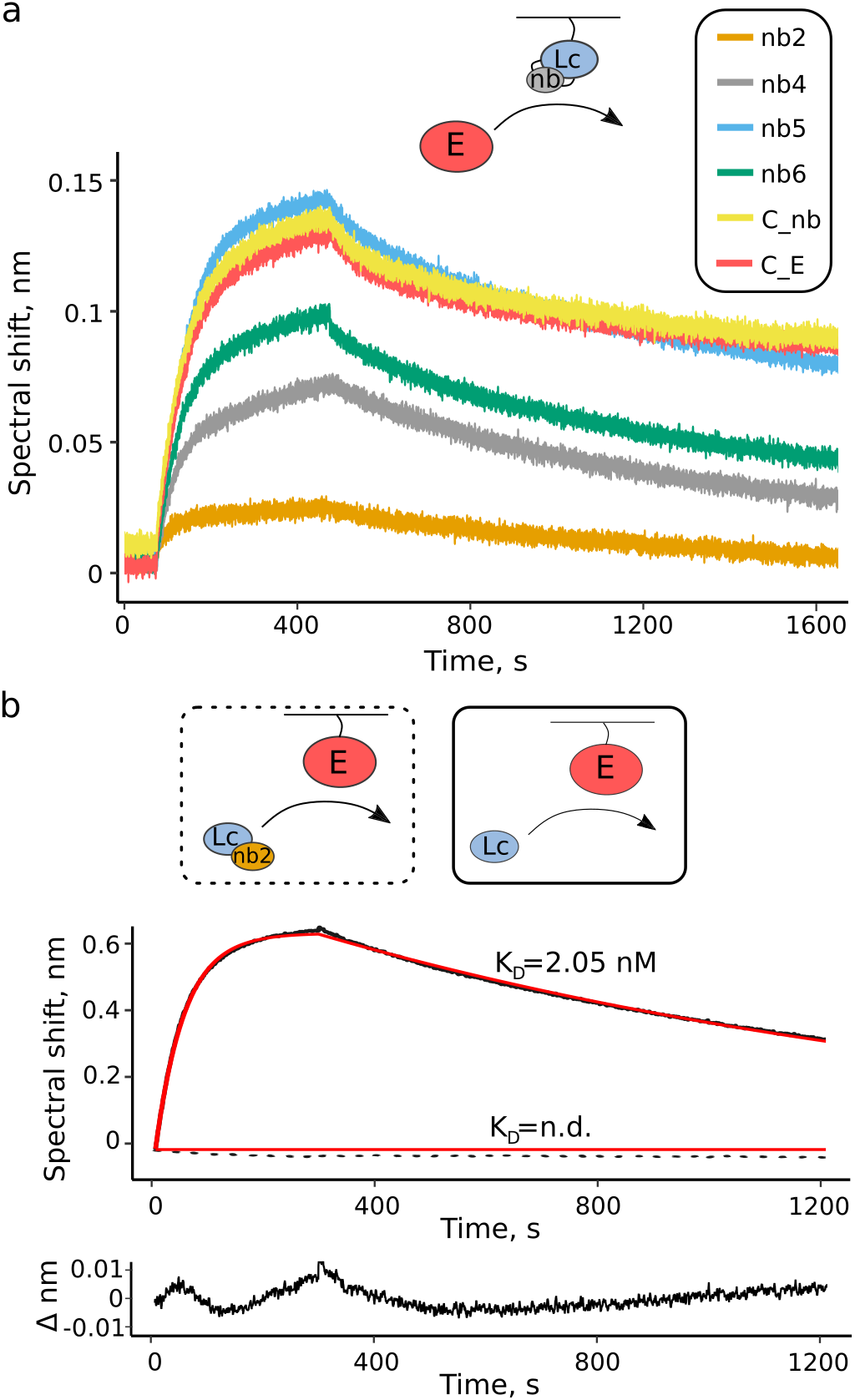
nb2 prevents GspL^cyto^ from binding the ATPase GspE. Upper panels contain schematic representation of each experiment and bottom panels contain the corresponding BLI traces, **(a)** Scouting for a GspL^cyto^ antagonist. Upon binding their epitopes, nanobodies 2, 4, 5 and 6 were crosslinked with GspL^cyto^ (symbolized by two linking lines on the scheme). The panel shows binding of GspE by GspL^cyto^-nanobody complexes. C_nb: negative control, GspE in the presence of an Flt3 receptor specific nanobody, C_E: negative control, GspE in the absence of a nanobody, **(b)** GspL^cyto^ is not capable to bind the ATPase upon binding nb2. BLI traces (in black; solid: GspL^cyto^, dashed: GspL^cyto^-nb2 as analytes, respectively) were fitted with 1:1 binding model (in red). The lowest panel shows the fit residuals. n.d. not determined.

For the experiments shown in **Figure 3** the GspE functionalized biosensors were exposed to different concentrations of an analyte (GspL^cyto^: 300-0.2 nM, GspLM: 300-0.2 nM). To verify that no aspecific binding was taking place, non-functionalized biosensors exposed in parallel to each ligand concentration as well as a functionalized biosensor dipped into kinetic buffer were employed as controls and the acquired data was used for subtraction during data processing. The GspE-GspL^cyto^ data were fitted with the ForteBio Data Analysis 7.1 software using a 1:1 interaction model.

For the experiments shown in **Figure 4** the nanobody functionalized biosensors were exposed to GspLM at 300 nM (panel a), and to GspL^peri^, GspM^peri^ and GspL^cyto^ at 500 nM each, sequentially (panel b). A non-functionalized biosensor immersed in analyte(s) solutions was used as a control.

For the experiment shown in **Figure 5a** the GspL^cyto^ functionalized biosensors were exposed to periplasmic fractions of nanobodies at ~1 μM followed by dipping in GspE at ~200 nM. A nonfunctionalized biosensor immersed in analyte(s) solutions was used as a control.

For the experiment shown in **Figure 5b** the GspE functionalized biosensors were in parallel exposed to (i) GspL^cyto^ at 57 nM (ii) nanobody 2 at 485 nM and GspL^cyto^ at 57 nM. In addition, the functionalized biosensors were exposed to (iii) kinetic buffer and (iv) nanobody 2 at 485 nM that served as reference biosensors.

### Nanobody generation and preparation

A library of single domain camelid antibodies, known as nanobodies (nb), was generated by immunizing a llama with an amphipol embedded GspLM (see paragraph: Membrane preparation and purification of recombinant GspLM in this section). The nanobodies were generated at the VIB Nanobody Service Facility [https://corefacilities.vib.be/nsf]. The generated library consists of 66 GspLM-specific nanobodies divided in eight CDR3 classes. In the used labels, the nanobody abbreviation nb is followed by a number that represents the class of the nanobody. Sequence alignment was performed for each class, and nanobodies having the canonical sequence were chosen as class representatives. The example of a (fragment of a) canonical sequence among four representatives is shown below in italics, whereas residues differing from the canonical ones are labeled in bold:

QVQLQESGGG**W**VQPG**N**SL**T**LSCAASGF**V**FS
QVQLQESGGGLVQPGGSLRLSCAASGF**D**FS
*QvQLQeSGGGLvQpGGSLRLSCAASGFFFS*
QVQLQESGGG**S**V**RT**GGSL**T**LSCAASGFFFS

The nanobodies were recloned to a modified pET15b vector, which introduced the N-terminal caspase-cleavable His-tag (see **Table S1**). The nanobody expression protocol is summarized in **Table S1** and described in a reference (51). For BLI experiments shown in **Figures 4** and **5a**, the periplasmic fraction containing nanobodies was used. For the experiment shown in **Figure 5b**, nanobodies were purified with two chromatographic steps (IMAC and SEC). The periplasmic fraction was prepared according to the published protocol, following the expression in minimal medium [54]. The purity and concentration of nanobodies enriched in the periplasmic fraction were checked on SDS-PAGE. Based on these estimations, the periplasmic fractions were diluted 10-40 times in the appropriate kinetic buffer prior to a BLI experiment.

## Author contributions

AF designed and performed recombinant protein production and biochemical and biophysical studies. IR purified and transferred to amphipols the GspLM complex used for llama immunizations. AF and SNS analyzed the data and wrote the manuscript with contributions from IR. SNS conceived and supervised the project.

## Acknowledgements

AF was supported by a predoctoral research fellowship from Research Foundation Flanders (FWO, Belgium). This project was supported by a GOA grant from Ghent University, core funding by the VIB, and an infrastructure grant AUGE-11–029 from the Hercules Foundation (Belgium) (to SNS).

## SUPPORTING INFORMATION

Additional supporting information may be found online in the Supporting Information section at the end of the article.

**Table S1.** Constructs used in the study.

**Table S2.** Affinities of GspL^cyto^-nanobody complexes.

**Figure S1.** Nanobodies, affiliation by complimentarity derermining regions.

**Supporting Figure S1.**
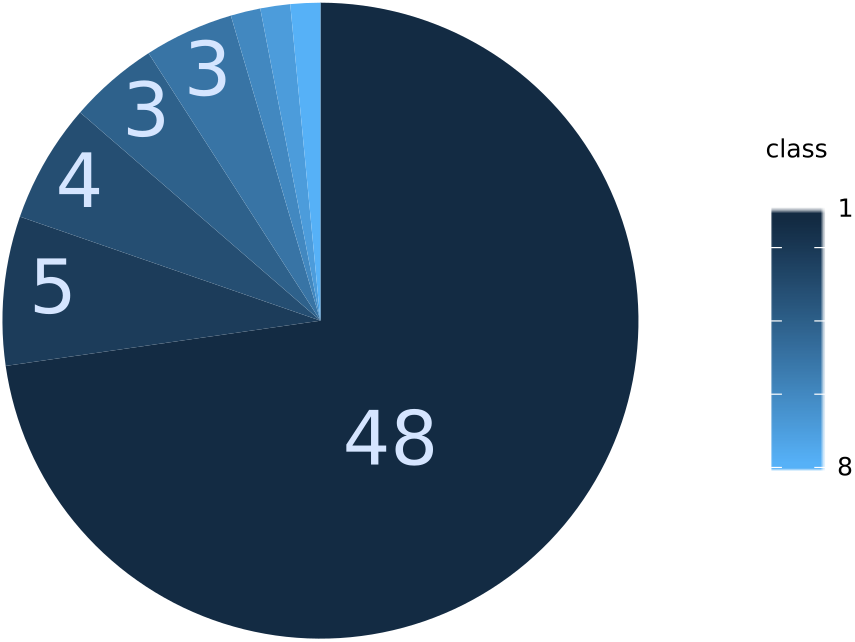
Nanobodies, affiliation by complementarity-determining regions. The generated library consists of 66 nanobodies belonging to eight different CDR3 classes. Each class is indicated with a ditterent shade of blue and size of a pie corresponds to the number of representatives, written in light blue. Classes 6, 7 and 8 have only single representatives.

**Supporting Table S2.** Affinities of GspL^cyto^ - nanobody complexes. The affinities were determined by BLI and are reported here along with the kinetic parameters of each interaction.

| nanobody | K<sub>D</sub> [M] | k<sub>on</sub> [1/Ms] | k<sub>off</sub> [1/s] |
| --- | --- | --- | --- |
| nb2 | 2.09 · 10<sup>-8</sup> | 9.14 · 10<sup>4</sup> | 1.91 · 10<sup>-3</sup> |
| nb4 | 2.24 · 10<sup>-8</sup> | 2.05 · 10<sup>5</sup> | 4.58 · 10<sup>-3</sup> |
| nb5 | 1.77 · 10<sup>-8</sup> | 7.63 · 10<sup>4</sup> | 1.35 · 10<sup>-4</sup> |
| nb6 | 3.32 · 10<sup>-8</sup> | 2.33 · 10<sup>5</sup> | 7.76 · 10<sup>-3</sup> |

